# Co-infection with two α-synuclein strains reveals novel synergistic interactions

**DOI:** 10.1101/2025.08.17.670736

**Authors:** Sara A. M. Holec, Chase R. Khedmatgozar, Shelbe J. Schure, Jason C. Bartz, Amanda L. Woerman

## Abstract

In synucleinopathies, the protein α-synuclein misfolds into Lewy bodies (LBs) in patients with Lewy body disease (LBD) or into glial cytoplasmic inclusions (GCIs) in patients with multiple system atrophy (MSA). The ability of a single misfolded protein to cause disparate diseases is explained by the prion strain hypothesis, which argues that protein conformation is a major determinant of disease. While structural, biochemical, and biological studies show that LBD and MSA patient samples contain distinct α-synuclein strains, we recently reported the unexpected finding of a novel α-synuclein strain in a Parkinson’s disease with dementia patient sample containing GCI-like co-pathology along with widespread LB pathology. This finding led us to question if two α-synuclein strains can interact with one another in a patient and, if so, can strain competition occur. Notably, this would not only impact the clinical presentation of disease but would also have profound impacts on successful therapeutic development. To test this possibility, we used the strain interference superinfection model developed in the prion field, in which a slower replicating strain—in this study, mouse-passaged MSA—is used to compete with a faster replicating strain—here, recombinant preformed fibrils (PFFs)— following sciatic nerve (sc.n.) inoculation. Unexpectedly, we found that PFFs generated using the same method differed in their ability to neuroinvade following sc.n. inoculation based on α-synuclein monomer source. Using a PFF preparation that does spread from the periphery, we conducted strain competition studies by first injecting TgM83^+/−^ mice with mouse-passaged MSA into the sc.n. followed by a second injection with PFFs at 30, 45, and 60% of the MSA incubation period. Unlike in the prion field, where the faster replicating strain inhibits the slower strain at the 30 and 45% time points, we found that the two α-synuclein strains exhibited a synergistic effect during neuroinvasion. Notably, disease onset across the three cohorts was shortened compared to MSA inoculation alone, and brains from terminal animals showed evidence of both the PFF and mouse-passaged MSA strains, suggesting the two strains worked together to accelerate neuroinvasion in the mice. These findings have important implications for disease progression in patients with α-synuclein co-pathologies. The finding that two strains can synergize with one another to accelerate the progression of clinical disease represents a novel outcome in mixed infection studies and more broadly expands our understanding of the effect of prion strain biology on disease pathogenesis.

## INTRODUCTION

Multiple system atrophy (MSA) and the Lewy body diseases (LBDs)−Parkinson’s disease, dementia with Lewy bodies, and Parkinson’s disease with dementia (PDD)−are part of a group of neurodegenerative diseases called synucleinopathies, which are defined by misfolding of the protein α-synuclein into pathological lesions in the brain (1–4). While these diseases are all linked to α-synuclein, each is discriminated by its unique clinical signs and affected cell types within the brain. This phenomenon is explained by the prion strain hypothesis, which proposes that clinical disease is determined by misfolded protein conformation (5–9). For example, in MSA, the pathological hallmark, glial cytoplasmic inclusions (GCIs), are localized primarily in oligodendrocytes, whereas in LBDs, the defining Lewy bodies (LBs) are found in neurons (1–4). While the differential presence of GCIs versus LBs is typically used to diagnose disease post-mortem, recent studies from our lab identified GCI-like co-pathology in the substantia nigra of a PDD patient sample (10). This unexpected finding led us to question how two distinct α-synuclein pathologies, or strains, interact with one another to impact clinical presentation and disease progression in synucleinopathy patients.

Research in the prion field has shown that multiple prion protein (PrP^Sc^) strains can exist in the same brain region in people with Creutzfeldt-Jakob disease (CJD) (11–13). A major consequence of these mixtures is strain competition, or the ability of one strain to interfere with or impede replication of a second strain. The ability of two strains to compete with one another was first shown following intracerebral (i.c.) inoculations of two PrP^Sc^ strains—the slower replicating strain 22C, and the faster replicating strain 22A—where replication of 22A in mice was blocked by prior inoculation with 22C, thereby extending the incubation time (14). Moreover, this same phenomenon was observed following extraneural intraperitoneal inoculation with the two strains, however, inactivation of the slower strain prior to injection allowed the faster strain to emerge and cause disease (15, 16). Subsequent studies in hamsters found that not only does staggering inoculation of two distinct strains result in the slower strain blocking replication of the faster strain, but also that the slower strain predominates in terminal animals following co-inoculation of the same two strains (17, 18). The suppression of a faster strain by a slower strain is now modeled using sciatic nerve (sc.n.) inoculations, forcing strains to replicate along the same neuroanatomical tracts during neuroinvasion, thus competing for the substrate needed to propagate. Notably, studies establishing this model revealed that the inhibitory effect seen with strain competition is dependent on the relative onset of replication of each strain; increasing the time between injections or decreasing the starting titer of each strain tested will prevent strain competition from occurring (18–20).

Combining our previous finding of GCI-like pathology in a PDD patient sample (10) with the known ability of PrP^Sc^ strains to interfere with one another, here we sought to determine the ability of two different α-synuclein strains to compete with one another for substrate following sc.n. inoculation. To accomplish this goal, we first identified our ‘fast strain’ by investigating the capacity of preformed fibrils (PFFs) made from recombinant α-synuclein containing the A53T mutation to neuroinvade from the periphery, given our previous findings that PFFs exhibit a shorter incubation period following i.c. inoculation in transgenic mice compared to MSA patient samples (21–23). Our prior work relied on PFFs generated using monomer purchased from either rPeptide (23) or Sigma (21, 22, 24). In comparing the two different monomer sources, we discovered that while both rPeptide and Sigma PFFs induced neurological disease following i.c. inoculation into the A53T-expressing TgM83^+/−^ mouse model, unexpectedly, only the rPeptide PFFs induced disease following sc.n. inoculation. We therefore used the rPeptide PFFs in strain competition studies with a mouse-passaged MSA patient sample, which served as our ‘slow strain’ in these studies. In this model, TgM83^+/−^ mice were first injected with the slow strain into the sc.n. prior to a second inoculation with the fast strain at time points 30, 45, and 60% of the slow strain incubation period. Surprisingly, we found that rather than suppression, the two strains worked synergistically to increase the rate of disease progression. Consistent with this outcome, we found evidence of both the PFF and MSA strains in the brains of terminal animals, suggesting the two strains colluded to exacerbate disease progression. All together, these data have important implications both for the use of PFFs in modeling synucleinopathies, as well as our understanding how co-pathologies of the same protein may synergize with one another to influence disease progression in human patients.

## MATERIALS AND METHODS

### Ethics statement

Animals were maintained in an animal facility in compliance with the 8th *Guide for the Care and Use of Laboratory Animals*. All procedures used in this study were approved by the University of Massachusetts Amherst and Colorado State University Institutional Animal Care and Use Committees.

### Generation of preformed fibrils (PFFs)

Recombinant human α-synuclein containing the A53T mutation was purchased from both Sigma and rPeptide. The lyophilized powder was resuspended in 1× DPBS to a concentration of 5 mg/mL and was fibrillized by incubating at 37 °C with constant agitation (1,200 rpm) in an orbital shaker for 7 d. PFFs were diluted with 1× DPBS to 1 mg/mL and stored at 4 °C until use.

### Cellular assay for α-synuclein prion quantification assay

Frozen brain tissue samples were used to generate 10% (wt/vol) brain homogenates in 1× DPBS and aggregated α-synuclein was isolated from these samples using sodium phosphotungstic acid (NaPTA; Sigma) precipitation, as previously reported (25), for use on cells constitutively expressing α-synuclein fusion proteins. The resulting protein pellets were diluted in 1× DPBS (1:14 for cells expressing the A53T mutation and 1:20 for cells expressing the V55Y mutation) before infecting HEK293T cells expressing mutant human α-synuclein fused to YFP, as described previously (26). The Sigma and rPeptide PFFs were diluted in 1× DPBS to 25 μg/mL before cell infection.

For all assays, samples were tested in 6 technical replicate wells of a 384-well plate covered with an adhesive membrane for 4 d at 37 °C. On day 4, DAPI and YFP images were collected from 4 regions of interest per well using the Lionheart FX automated fluorescent microscope. BioTek Gen5 software was used to build custom algorithms to quantify the total fluorescence summed across all aggregates normalized to cell count (× 10^5^, arbitrary units, A.U.). Values from the 4 regions of interest were combined to determine a single value for each well, and the 6 replicate well values were then averaged to calculate prion infectivity for a single sample.

### Mice

All mice were group housed, unless health concerns deemed separation necessary. Animals were maintained in ABSL-2 conditions with free access to water and food on a 12-hr light/dark cycle. The TgM83^+/−^ mice used here were generated by breeding TgM83^+/+^ male mice (27) with B6C3F1 female mice, both of which were purchased from The Jackson Laboratory.

### Mouse inoculations

Frozen mouse brain tissue from TgM83^+/−^ mice inoculated with either a control or an MSA patient sample was homogenized in 1× Dulbecco’s phosphate buffered saline (DPBS) using the Omni Tissue Homogenizer (Omni International) to generate a 10% (wt/vol) homogenate. The resulting homogenate was then diluted to 5 mg/mL in 1× DPBS. For intracranial (i.c.) inoculations, 6-to-7-week-old TgM83^+/−^ mice were anesthetized via isoflurane prior to using a freehand injection to deliver 20 μL of 1 mg/mL PFFs or 5 mg/mL brain homogenate into the thalamus. For sciatic nerve (sc.n.) inoculations, 12-week-old TgM83^+/−^ mice were anesthetized via isoflurane prior to injecting 2 μL of 1 mg/mL PFFs or 5 mg/mL brain homogenate into the sc.n., as previously described (23). Briefly, hair was removed before making a small incision (≤1 cm) in the left hindlimb to expose the sc.n. A 28-gauge needle was moved up and down parallel to the nerve prior to the injection. The incision was closed with vicryl rapide suture.

Two sc.n. inoculations were performed for superinfection studies using the procedure described above. The first inoculation was performed using 20 μL of 5 mg/mL mouse-passaged MSA patient sample in 12-week-old TgM83^+/−^ mice. The second inoculation was performed at 30%, 45%, and 60% of the previously reported incubation period for MSA inoculation via sc.n. (23). These inoculations correspond to 63 days post-inoculation (dpi), 92 dpi, and 126 dpi, respectively. The second inoculation was performed using 20 μL of 1 mg/mL PFFs using monomer from rPeptide.

After recovery, mice were evaluated 3 times per week for the onset of neurological signs using standard prion disease criteria (28). Mice inoculated with PFFs, mouse-passaged MSA patient sample, or superinfection with both strains were euthanized following the onset of progressive neurological signs. Mice inoculated with passaged control patient sample were euthanized at 365 dpi following i.c. inoculation, or 500 dpi following sc.n. inoculation. Following euthanasia, the brain was removed and bisected down the midline. One half was frozen for subsequent biological analysis and the other half was formalin-fixed for neuropathological assessment. Additionally, the spinal column from mice inoculated with PFFs into the sc.n. was also collected in formalin to assess neuropathology.

### Immunohistochemistry and neuropathology

The formalin-fixed half-brains were coronally sectioned prior to processing through graded alcohols, xylene, and paraffin, followed by embedding. The embedded brain tissue was cut on a microtome into 8 μm sections, mounted, and deparaffinized. Fixed spinal columns were decalcified in Rapid Bone Decalcifier (StatLab) and cut into 5 mm sections prior to processing as described above. The spinal column was then cut using a microtome into 10 μm sections, mounted, and deparaffinized. Heat-mediated antigen retrieval was performed using citrate buffer (0.1 *M*, pH 6) for 20 min. All slides were incubated at room temperature overnight in primary antibodies for α-synuclein phosphorylated at S129 (EP1536Y; 1:1,000; Abcam) and glial fibrillary acidic protein (GFAP; 1:500; Abcam) after blocking in 10% (vol/vol) normal goat serum. Secondary antibodies conjugated to AlexaFluor 488 and 594 (1:500; ThermoFisher) were used to detect primary antibodies. Nuclei were labelled with Hoechst 33342 (1:5,000;

ThermoFisher). Slides were imaged using the Lionheart FX automated microscope (Agilent BioTek) with the 10x objective. Quantification of neuropathology was performed using the BioTek Gen5 software package. Pathology was measured by setting a pixel intensity threshold using both positive and negative control slides. Regions of interest were drawn around the caudoputamen (Cd), piriform cortex and amygdala (Pir), hippocampus (HC), thalamus (Thal), hypothalamus (HTH), midbrain (Mid), and pons. The positive pixel percentage in each region was averaged across all mice in each experimental group. Representative images of the brain and spinal cord were acquired using the Leica DMi8 inverted fluorescent microscope and processed using LAS X software.

### Statistical analysis

All data are presented as mean ± standard deviation. Data were analyzed using GraphPad Prism software (version 10). Cell assay data comparing the passaged control and PFF inocula (both Sigma and rPeptide) were analyzed using a one-way analysis of variance (ANOVA) test with a Tukey’s multiple comparison post hoc test. Kaplan-Meier curves were analyzed using a log-rank Mantel-Cox test. Neuropathology data were analyzed using a two-way ANOVA with a Tukey’s multiple comparison post hoc test.

Cell assay data comparing mouse samples from animals inoculated i.c. and sc.n. using PFFs were analyzed using a two-way ANOVA with a Tukey’s multiple comparison post hoc test. Cell assay data comparing superinfection samples were analyzed using a one-way ANOVA with a Tukey’s multiple comparison post hoc test. Significance was determined with a *P*-value < 0.05.

## RESULTS

### Infectivity of α-synuclein PFFs in α-syn140-YFP bioreporter cells is impacted by monomer source

The concept of strain competition was previously investigated using multiple PrP prion strains, with studies showing that strain interference results in the ability of a faster replicating prion strain to inhibit the spread of a second slower replicating strain (14, 17–19, 29–31). To investigate whether or not α-synuclein strains can compete with one another, we selected two α-synuclein fibril sources previously used by our group with distinct incubation periods — recombinant A53T PFFs and a TgM83^+/−^-passaged MSA patient sample (23). Using our panel of α-syn140-YFP bioreporter cell lines (26) to define the biological properties of each strain, we recently showed that while several strains, including MSA, replicate in α-syn*A53T-YFP cells, our α-syn*V55Y-YFP cell line can differentiate between MSA and A53T PFFs following mouse passage based on selective replication of the passaged PFFs *in vitro* (23). These recent studies used PFFs that were generated using monomer sourced from rPeptide, however, our prior studies investigating α-synuclein strain biology used monomer purchased from Sigma, instead (21, 22, 24). To determine if the source of α-synuclein monomer impacts the resultant PFF biology, thereby impacting data from our strain interference studies, we first tested the ability of both Sigma and rPeptide A53T PFFs to replicate in the α-syn*A53T-YFP and α-syn*V55Y-YFP cells (Figure 1). Interestingly, we found that the rPeptide PFFs were more infectious in both the A53T (66 ± 10 × 10^5^ A.U.) and V55Y cells (45 ± 7.6 × 10^5^ A.U.) compared to the Sigma PFFs (A53T: 23 ± 4.8 × 10^5^ A.U.; V55Y: 12 ± 2.9 × 10^5^ A.U.; *P* < 0.0001). However, the Sigma PFFs still induced significant infection in both cell lines compared to the negative control (A53T: 2.0 ± 0.4 × 10^5^ A.U.; V55Y: 2.1 ± 1.1 × 10^5^ A.U.; P ≤ 0.0001). The negative control data, generated using TgM83^+/−^-passaged negative control patient sample, was previously reported (23). Overall, as our bioreporter cells are predictive of sample titer (32, 33), these data suggest potential underlying differences in the biological activity between the two monomer sources.

**Figure 1.**
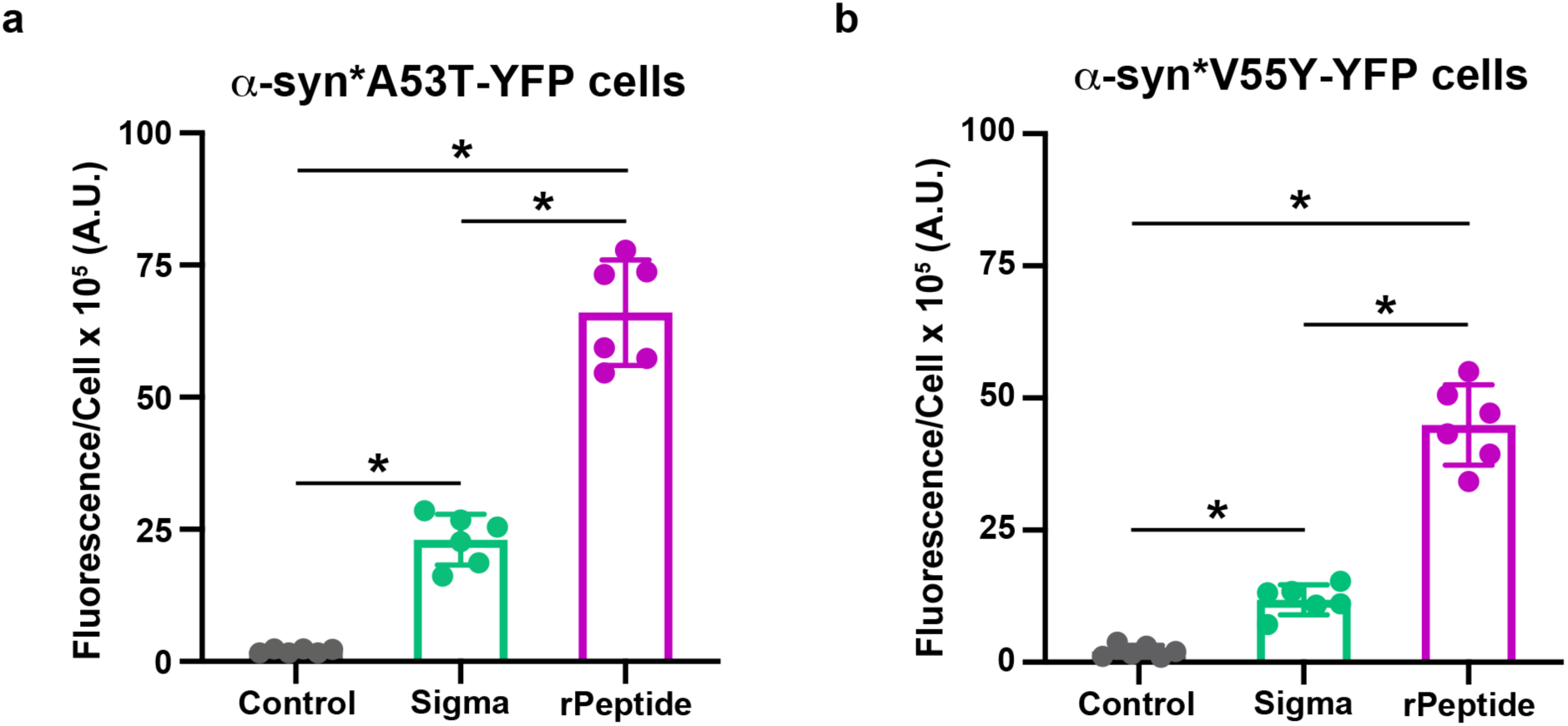
Monomer source used to generate preformed fibrils impacts infectivity in α-syn140-YFP cells. Monomeric A53T α-synuclein purchased from either Sigma (green) or rPeptide (purple) was fibrillized in 1× DPBS with shaking for 1 week to generate preformed fibrils (PFFs). The resulting fibrils were tested for infectivity in the (a) α-syn*A53T-YFP and (b) α-syn*V55Y-YFP cell lines. Compared to mouse-passaged control patient sample (gray), both the Sigma and rPeptide A53T PFFs induced significant infection in both cell lines. Quantification of α-synuclein prion infectivity (× 10^5^ arbitrary units [A.U.]) shown as mean ± standard deviation. Negative control data previously published (23). * = *P* < 0.05.

### Disease pathogenesis of α-synuclein PFFs in TgM83^+/−^ mice is impacted by protein source

We next compared the ability of the two different PFFs to transmit disease following i.c. or sc.n. inoculation into the TgM83^+/−^ mouse model (Figure 2). Consistent with the stronger cellular infection, we found that i.c. injection using the rPeptide PFFs induced disease in 78 ± 6 days post-inoculation (dpi), whereas disease onset using the Sigma PFFs was delayed until 107 ± 22 dpi (*P* < 0.05; Figure 2a). Both i.c. cohorts resulted in a 100% attack rate, inducing disease in all 7 of the injected mice. Moreover, both PFF sources were significantly different compared to mice inoculated with a mouse-passaged control patient sample, which remained asymptomatic through 380 dpi (*P* > 0.001; 0% attack rate in 10 mice). The control and rPeptide i.c. inoculation data were previously published elsewhere (23). Unexpectedly, when we used the same PFFs to inoculate TgM83^+/−^ mice into the sc.n., the rPeptide PFFs induced disease in 112 ± 8 dpi in 100% of the 7 mice injected, but mice inoculated with the Sigma PFFs remained asymptomatic through 475 dpi (Figure 2b; *P* < 0.0001; 0% attack rate in 10 mice). These data suggest that the two monomer sources impact protein misfolding into distinct conformations during fibrillization, with the Sigma PFFs unable to efficiently replicate in the peripheral nervous system. Consistent with this interpretation, the Sigma PFF cohort was indistinguishable from mice inoculated with the negative control sample, which were also asymptomatic through 475 dpi [data previously published (23); 0% attack rate in 10 mice].

**Figure 2.**
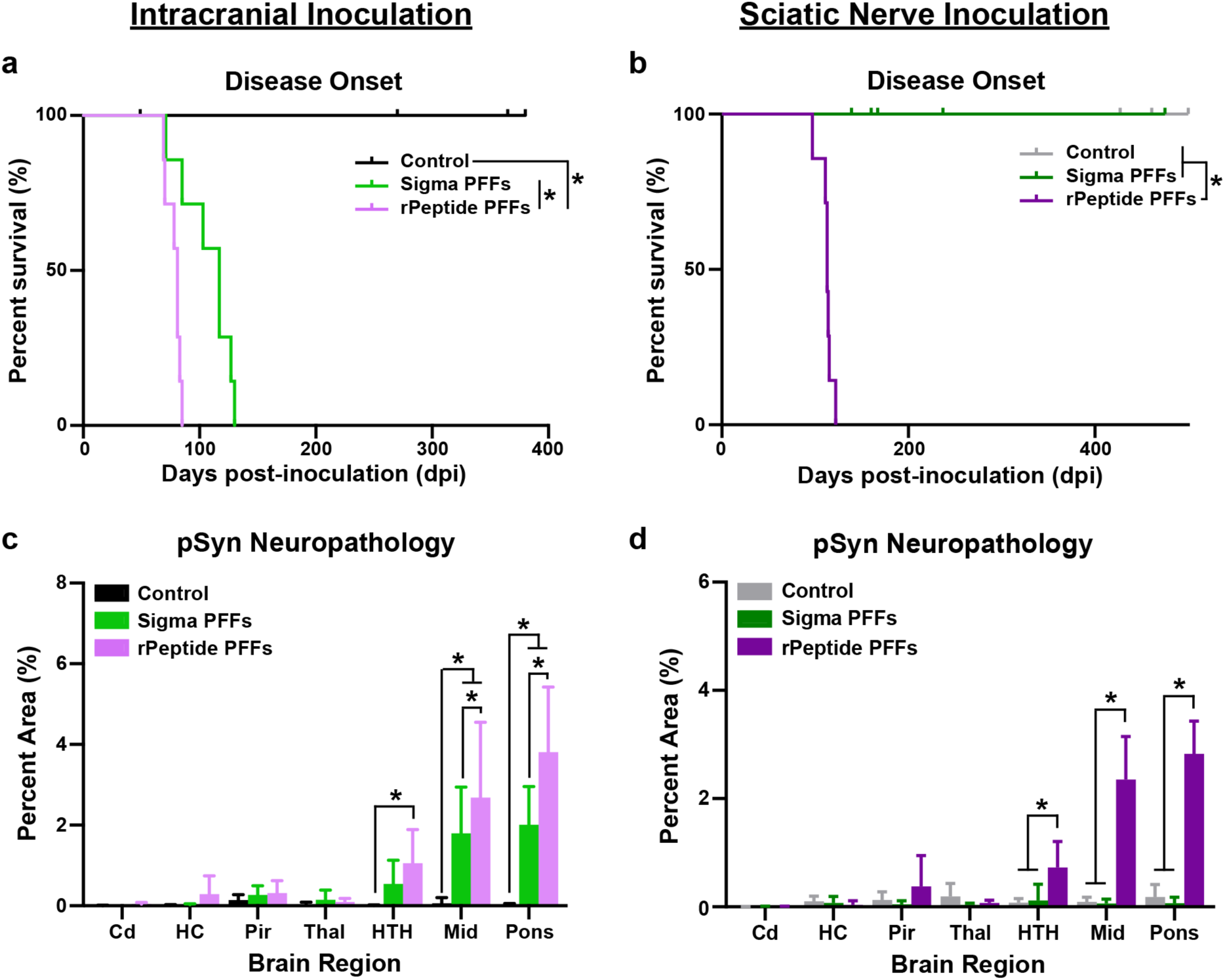
Monomer source used to generated preformed fibrils impacts disease transmission to TgM83^+/−^ mice. Monomeric A53T α-synuclein purchased from either Sigma (greens) or rPeptide (purples) was fibrillized in 1× DPBS with shaking for 1 week to generate preformed fibrils (PFFs). PFFs were diluted to 1 mg/mL and mouse-passaged negative control sample (black and gray) was diluted to 5 mg/mL before 20 μL were then inoculated either (a & c) intracranially (i.c.) or (b & d) into the sciatic nerve (sc.n.) in TgM83^+/−^ mice. (a & b) Kaplan-Meier plot showing disease onset in TgM83^+/−^ mice inoculated (a) i.c. or (b) sc.n. Data for both i.c.- and sc.n.-inoculated control mice, as well as mice inoculated i.c. with rPeptide A53T PFFs were previously reported (23). (c & d) Quantification of phosphorylated α-synuclein neuropathology (EP1536Y primary antibody, 1:1,000 dilution) in the caudate (Cd), hippocampus (HC), piriform cortex and amygdala (Pir), thalamus (Thal), hypothalamus (HTH), midbrain (Mid), and pons of TgM83^+/−^ mice following (c) i.c. or (d) sc.n. inoculation with mouse-passaged control, Sigma A53T PFFs, or rPeptide A53T PFFs. Neuropathology from control (i.c. and sc.n.) and rPeptide A53T PFF (i.c.) inoculated mice were previously reported (23). Data shown as mean ± standard deviation. * = *P* < 0.05.

Following the onset of neurological signs or experimental endpoint, dissected brains were bisected along the midline with one half fixed in formalin for neuropathological analysis. The resulting α-synuclein lesions were stained using an antibody specific to α-synuclein phosphorylated at S129 (EP1536Y), and neuropathology was quantified in the caudate (Cd), hippocampus (HC), piriform cortex and amygdala (Pir), hypothalamus (HTH), thalamus (Thal), midbrain (Mid), and pons. Terminal mice inoculated i.c. with either PFF source developed significant inclusions in the Mid and pons compared to the control-inoculated mice (Figure 2c; *P* < 0.0001). Notably, the rPeptide PFFs induced more neuropathology in the Mid and pons than the Sigma PFFs, as well as showed a significant increase in HTH inclusions compared to negative control (*P* < 0.05). Representative images of Sigma and rPeptide PFF-induced pathology following i.c. inoculation are shown in Figure 3a & b. Neuropathology data from mice i.c.-inoculated with control and rPeptide PFFs were previously reported (23). Consistent with the presence or absence of neurological signs, sc.n. inoculation of rPeptide PFFs induced α-synuclein pathology whereas the Sigma PFFs failed to induce detectable inclusions (Figure 2d). While changes were not detected in control-inoculated mice, the rPeptide-induced neuropathology was significantly increased in the HTH, Mid, and pons compared to both the control- and Sigma PFF-inoculated animals (*P* < 0.001). Representative images of neuropathology following sc.n. inoculation using the Sigma and rPeptide PFFs are shown in Figure 3c & d, respectively. Quantification of sc.n.-inoculated control sample was previously published (23).

**Figure 3.**
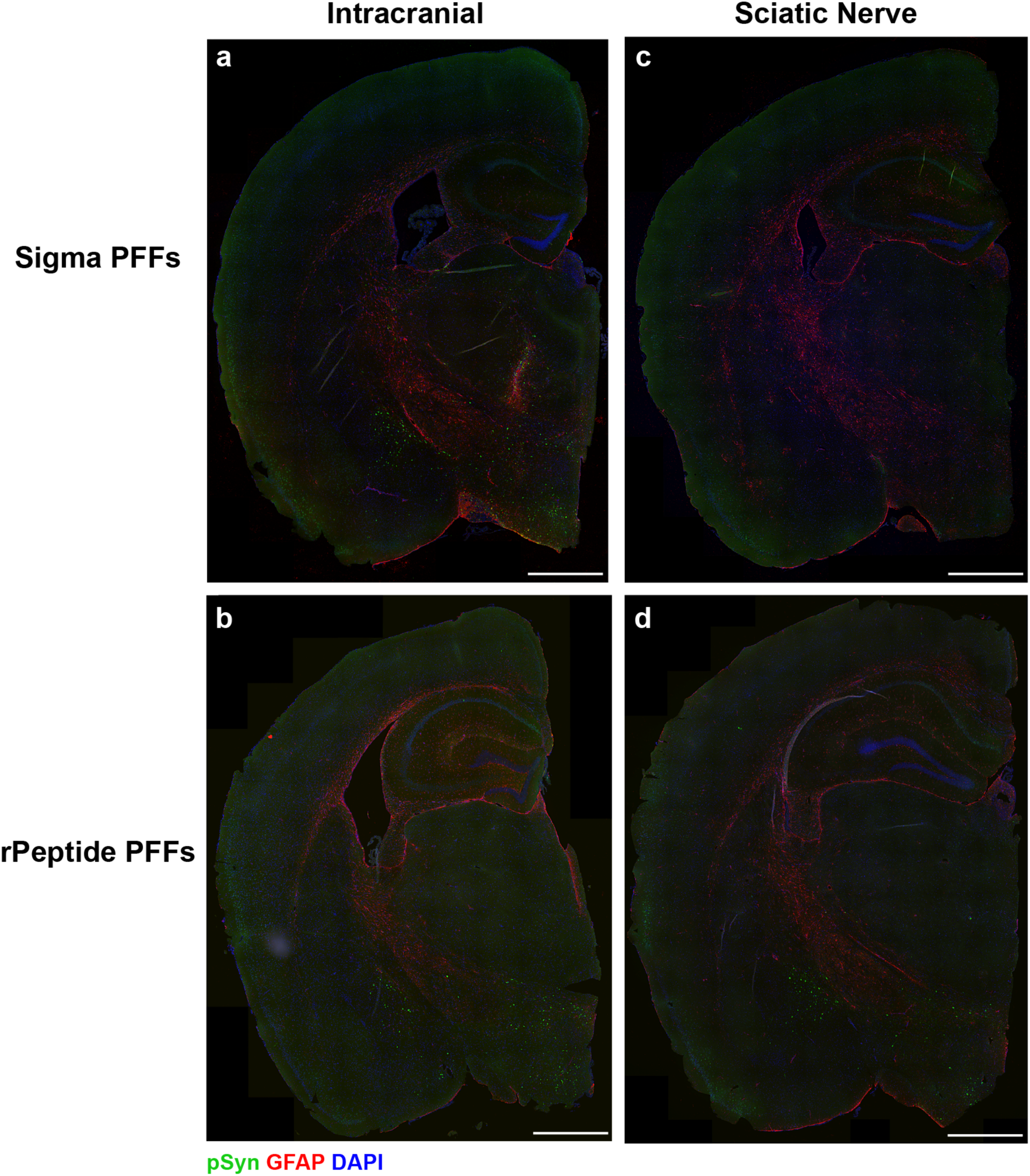
Sigma PFFs fail to induce α-synuclein neuropathology in the brain following sciatic nerve inoculation. Monomeric A53T α-synuclein purchased from either Sigma or rPeptide was fibrillized in 1× DPBS with shaking for 1 week to generate preformed fibrils (PFFs). PFFs were diluted to 1 mg/mL before 20 μL was then inoculated either (a & b) intracranially (i.c.) or (c & d) into the sciatic nerve (sc.n.) in TgM83^+/−^ mice. Sections including the hippocampus, piriform cortex, amygdala, thalamus, and hypothalamus are shown. Representative α-synuclein neuropathology (EP1536Y primary antibody; 1:1,000) in green, glial fibrillary acidic protein (GFAP; 1:500) in red, and DAPI in blue. Scale bar, 200 μm.

To determine if the lack of neurological disease and brain neuropathology following sc.n.-inoculation of Sigma PFFs was due to either failed neuroinvasion or slow replication resulting in sub-clinical disease, we stained fixed spinal cords using the same pS129 antibody (Figure 4). Sections along the full length of the spinal cord from mice inoculated with Sigma PFFs were negative for α-synuclein inclusions (Figures 4a & b) whereas the terminal rPeptide PFF-inoculated mice developed robust neuropathology (Figures 4c & d). These findings point to an inability of the Sigma PFFs to replicate in the periphery, whereas peripheral replication of the rPeptide PFFs enabled spread of disease into the central nervous system.

**Figure 4.**
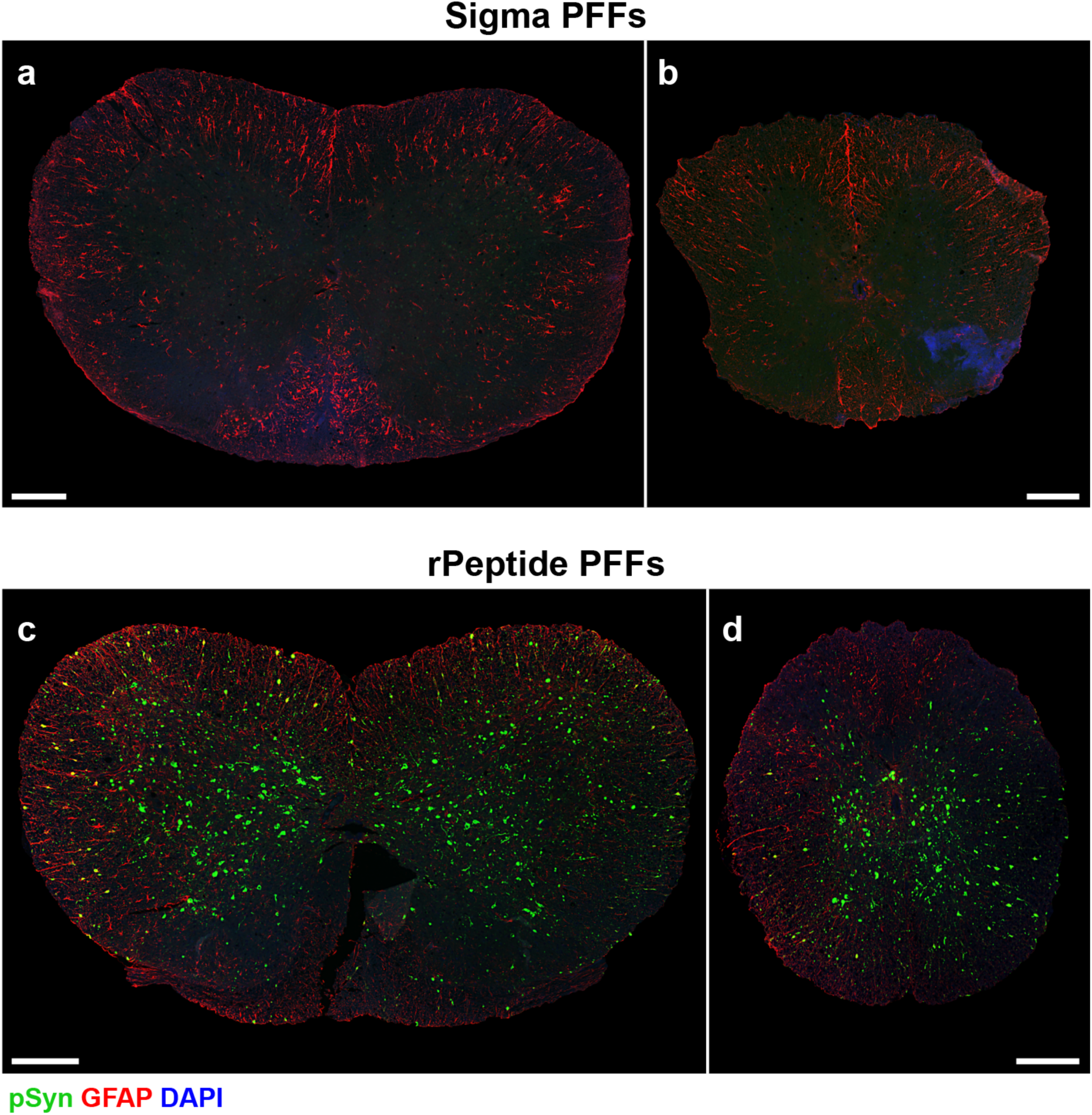
rPeptide PFFs induce spinal cord neuropathology following sciatic nerve inoculation into TgM83^+/−^ mice while Sigma PFFs do not. Monomeric A53T α- synuclein purchased from either Sigma or rPeptide was fibrillized in 1× DPBS with shaking for 1 week to generate preformed fibrils (PFFs). PFFs were diluted to 1 mg/mL before 20 μL was then inoculated into the sciatic nerve of TgM83^+/−^ mice. Representative images of the cervical (a & c) and thoracic (b & d) spinal cord after inoculation with (a & b) Sigma A53T PFFs or (c & d) rPeptide A53T PFFs. Phosphorylated α-synuclein (EP1536Y primary antibody; 1:1,000) in green, glial fibrillary acidic protein (GFAP; 1:500) in red, and DAPI in blue. Scale bar, 200 μm.

Finally, we isolated α-synuclein prions from frozen half brains collected from TgM83^+/−^ mice inoculated either i.c. or into the sc.n. and tested their ability to replicate in the α-syn*A53T-YFP and α-syn*V55Y-YFP cell lines (Figure 5). We found that samples from mice inoculated i.c. with the mouse-passaged negative control sample did not effect either cell line (A53T: 2.71 ± 1.34 × 10^5^ A.U; V55Y: 2.9 ± 1.1 × 10^5^ A.U.). By comparison, both the Sigma and rPeptide PFFs inoculated i.c. replicated in the α- syn140*A53T-YFP cells (Sigma: 13 ± 8.3 × 10^5^ A.U.; rPeptide: 13 ± 4.9 × 10^5^ A.U.; *P* < 0.01) and the α-syn140*V55Y-YFP cells (Sigma: 8.7 ± 5.1 × 10^5^ A.U.; rPeptide: 10 ± 5.6 × 10^5^ A.U.; *P* > 0.05). In comparison with the starting inocula (Figure 1), the mouse-passaged PFFs showed similar infectivity in both cell lines (*P* > 0.05). Consistent with our clinical and neuropathological observations, samples from mice inoculated with rPeptide PFFs into the sc.n. induced infection in both the α-syn140*A53T-YFP (20 ± 11 × 10^5^ A.U.) and α-syn140*V55Y-YFP cells (13 ± 5.2 × 10^5^ A.U.; *P* < 0.0001) whereas the Sigma PFF-inoculated mouse samples had no effect in either cell line (A53T: 4.4 ± 1.8 × 10^5^ A.U.; V55Y: 1.2 ± 0.4 × 10^5^ A.U.; *P* > 0.05). As expected, the control sc.n.-inoculated mouse samples were also negative for α-synuclein prion replication in the two cell lines (A53T: 1.8 ± 0.7 × 10^5^ A.U; V55Y: 2.1 ± 0.63 × 10^5^ A.U.). These data confirm the absence of sub-clinical replication of Sigma PFFs following sc.n. inoculation in TgM83^+/−^ mice. Moreover, these findings also indicate that the PFFs generated using the two different monomer sources misfold into at least two distinct conformations defined by a unique replication capacity and disease pathogenesis.

**Figure 5.**
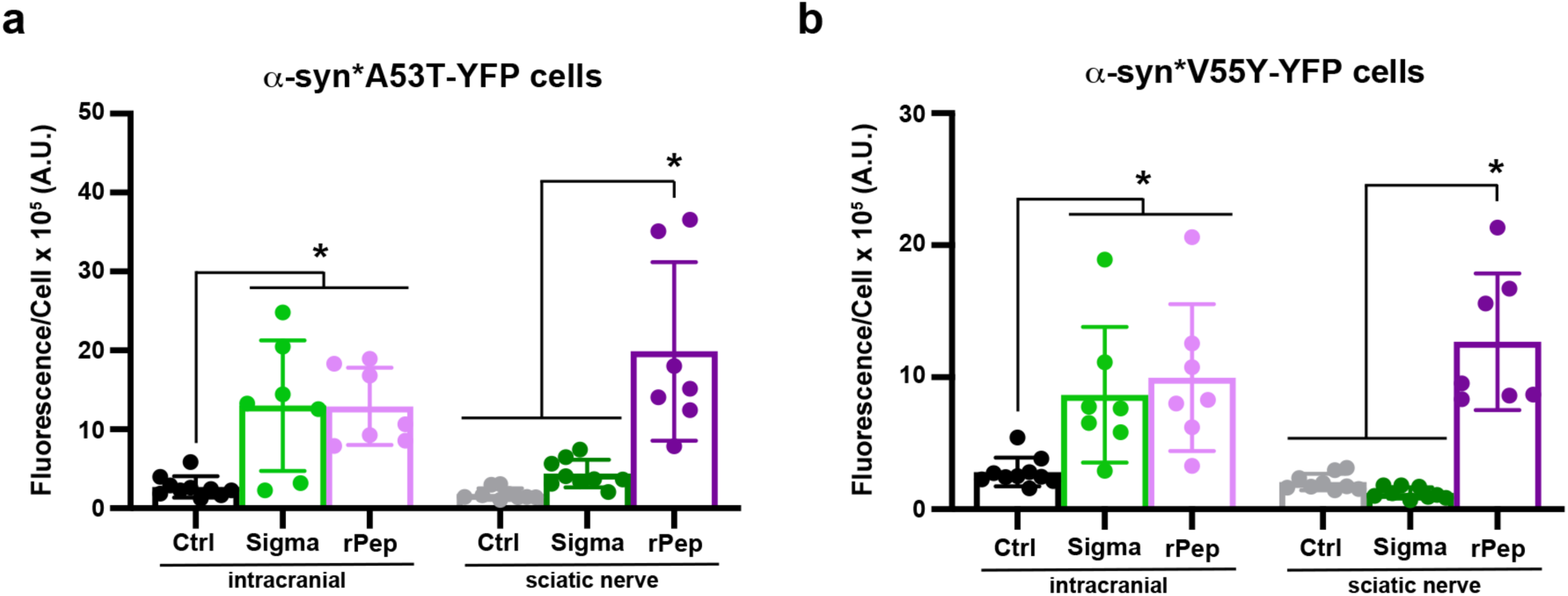
Different monomer sources generate A53T preformed fibrils with altered replication capacity. Monomeric A53T α-synuclein purchased from either Sigma (greens) or rPeptide (purples) was fibrillized in 1× DPBS with shaking for 1 week to generate preformed fibrils (PFFs). PFFs were diluted to 1 mg/mL and mouse-passaged negative control sample (black and gray) was diluted to 5 mg/mL before 20 μL were inoculated either intracranially (i.c.) or into the sciatic nerve (sc.n.) in TgM83^+/−^ mice. Frozen half-brains were homogenized in in 1× DPBS before isolating α-synclein prions via phosphotungstic acid precipitation. The isolated samples were tested for infectivity in the (a) α-syn*A53T-YFP and (b) α-syn*V55Y-YFP cell lines. PFFs inoculated i.c. induced α-synuclein prion formation capable of replicating in both cells lines, regardless of monomer source, whereas the negative control inoculations did not. By comparison, only the rPeptide PFFs induced α-synuclein prion formation following sc.n. inoculation. Samples from mice inoculated with Sigma PFFs or mouse-passaged negative control were unable to replicate in either cell line. Quantification of α-synuclein prion infectivity (× 10^5^ arbitrary units [A.U.]) shown as mean ± standard deviation. * = *P* < 0.05.

### Superinfection of mouse-passaged MSA and PFFs reveals a synergistic effect in onset of clinical disease

Having determined that Sigma PFFs are unable to neuroinvade following sc.n.-inoculation, we chose to use the rPeptide PFFs in comparison with mouse-passaged MSA in our strain competition studies. Superinfection models in the prion field rely on a model in which the animal is first challenged by the slower replicating strain via sc.n. inoculation followed by challenge with the faster replicating strain at 30, 45, or 60% of the slower strain incubation period (19, 20). This approach forces the two strains to replicate in the same cells, requiring competition with one another for available substrate (31).

Applying this same approach, we performed superinfection studies via injection into the sc.n. with mouse-passaged MSA into TgM83^+/−^ mice. A subsequent inoculation of rPeptide A53T PFFs was then performed at 30% (63 dpi), 45% (92 dpi), or 60% (126 dpi) of the previously reported mouse-passaged MSA incubation period (23) (Table 1). In PrP studies, the faster strain typically outcompetes the slower strain in the 30% cohort, with no effect on incubation period. Here, however, we found that our 30% superinfection cohort exhibited a slight increase in disease onset from 112 ± 8 dpi for the PFFs alone compared to 138 ± 18 dpi for the 30% superinfection, with a 100% attack rate in 9 mice (Figure 6a). We additionally found that the 30% superinfection cohort differed significantly from the 45% and 60% superinfection cohorts that had overlapping incubation periods of 173 ± 45 dpi and 176 ± 31, respectively (Figure 6a). In the 45% cohort, all 11 mice developed clinical disease. However, in the 60% cohort, 8 of the 11 mice injected developed neurological signs. The 3 additional mice died for non-experimental reasons between 90-115 dpi. Notably, all superinfection cohort incubation periods were shorter than the mouse-passaged MSA incubation period of 200 ± 49 dpi (Figure 6a). Neuropathological assessment of mouse samples across all three superinfection cohorts showed robust inclusions in the Mid and pons, demonstrating the presence of pathogenic α-synuclein following the peripheral challenge (Figure 6b).

**Fig. 6.**
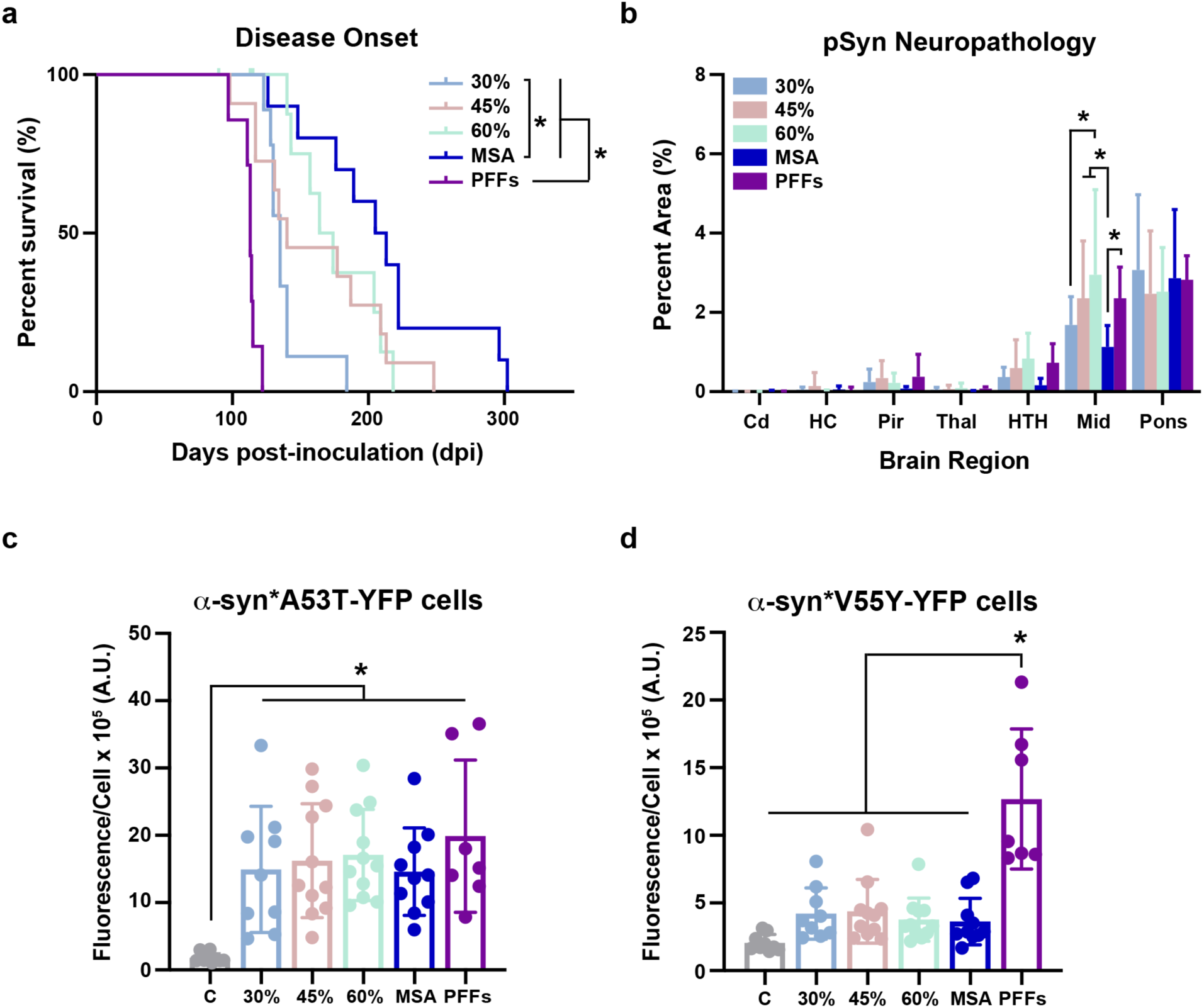
Superinfection of two α-synuclein strains leads to same disease outcome. (a) Kaplan-Meier plot showing disease onset in TgM83^+/−^ mice sc.n. inoculated with mouse-passaged MSA (blue), rPeptide A53T PFFs (purple), 30% superinfection (light blue), 45% superinfection (light pink), 60% superinfection (light green). (b) Quantification of phosphorylated α-synuclein neuropathology (EP1536Y primary antibody, 1:1,000 dilution) in the caudate (Cd), hippocampus (HC), piriform cortex and amygdala (Pir), thalamus (Thal), hypothalamus (HTH), midbrain (Mid), and pons of TgM83^+/−^ mice following sc.n. inoculation. PTA precipitation was performed on brain homogenates and samples were tested for infectivity in the (c) α-syn*A53T-YFP and (d) α-syn*V55Y-YFP cell lines. Quantification of α-synuclein prion infectivity (× 10^5^ arbitrary units [A.U.]) shown as mean ± standard deviation. * = *P* < 0.05.

**Table 1.** Superinfection incubation period in TgM83^+/−^ mice.

| Cohort | First inoculum | Second inoculum | Interval between inoculations | Onset of clinical signs |  | n/n<sub>0</sub><sup>b</sup> |
| --- | --- | --- | --- | --- | --- | --- |
|  |  |  |  | After 1<sup>st</sup> inoculation<sup>a</sup> | After 2<sup>nd</sup> inoculation<sup>a</sup> |  |
| **MSA** | MSA | — | n.a. | 210 ± 56 | n.a. | 10/10 |
| **PFFs** | — | PFFs | n.a. | n.a. | 112 ± 8 | 7/7 |
| **30%** | MSA | PFFs | 63 days | 138 ± 18 | 75 ± 18 | 9/9 |
| **45%** | MSA | PFFs | 92 days | 161 ± 48 | 69 ± 48 | 11/11 |
| **60%** | MSA | PFFs | 126 days | 157 ± 42 | 50 ± 31 | 11/11 |

<sup>a</sup>Days post-inoculation ± standard deviation; <sup>b</sup>Number of symptomatic animals (n) / number of mice inoculated (n<sub>0</sub>).

To determine which of the two α-synuclein strains was present in the brain at the time of clinical onset of disease, we assessed the biological activity of pathogenic α-synuclein present in each brain. We previously showed that terminal TgM83^+/−^ mice inoculated with MSA patient samples contain a consistent titer of α-synuclein prion in the brain (33). In line with these data, we found that brain tissue from all three superinfection cohorts showed similar infectivity in the α-syn140*A53T-YFP cells as the brains from mice inoculated with either A53T PFFs or mouse-passaged MSA (Figure 6c). These data indicate that similar amounts of pathogenic α-synuclein are present in the brain at onset of terminal disease, independent of strain. We then tested the same samples in α-syn140*V55Y-YFP cells (Figure 6d), as this mutation was previously shown to support replication of mouse-passaged A53T PFFs but not this specific mouse-passaged MSA patient sample (23). We found that none of the superinfection cohorts infected the α-syn140*V55Y-YFP cells, suggesting that an inhibitory dose of mouse-passaged MSA was present in the brain at the onset of terminal disease. Combining the incubation time and cell assay data, these findings suggest that the mouse-passaged MSA and PFF strains act synergistically, or collude with one another, to impact disease pathogenesis and increase the rate of neuroinvasion.

## DISCUSSION

The presence of co-pathologies in the brains of patients with neurodegenerative disorders has been recognized since the 1980s, when the pathologies first described by Alois Alzheimer were shown to include the β protein in amyloid plaques (34, 35) and the protein tau in neurofibrillary tangles (36, 37). Despite this awareness, the development of research models that enable scientists to interrogate how two or more proteins interact to cause or alter disease has only recently become a priority in the field (38–40). As an increasing number of animal models are developed with the objective of enabling research on co-pathologies [reviewed in (41)], this line of investigation has continued to ignore the effect co-pathologies of the same protein can have on disease progression. In contrast, over twenty years of research in the prion field has shown that CJD patients can develop multiple PrP^Sc^ strains, which likely interact to influence the clinical presentation of disease (11–13). Consistent with these findings, we recently described the presence of GCI-like inclusions in the brain of a patient diagnosed with PDD, who also had widespread LB pathology (10). This finding, which is not unique to this particular Lewy body disease patient (42), led us to question if and/or how two α-synuclein strains can interact with one another to alter disease progression.

Similar questions in the prion field resulted in models of strain competition, in which a slower replicating prion strain is used to block or inhibit the propagation of a faster strain as the two are forced to compete with one another for access to substrate (14–20). Notably, achieving this competition requires forcing replication to occur within the same cell population. This is achieved via sc.n. injection, rather than the more commonly used i.c. inoculations, as the two strains must spread from the periphery to the brain via the same neuroanatomical tracts (43). Applying this well-established approach here, we used a mouse-passaged MSA patient sample as our slow strain, which we previously analyzed for its ability to neuroinvade following sc.n. injection (23). In comparison, drawing on the use of recombinant α-synuclein PFFs as a model of synucleinopathy over the past 15 years (44–46), we chose to use PFFs harboring the A53T mutation as our faster replicating strain.

While it is now well-documented that PFFs fibrillized under varying conditions adopt distinct misfolded conformations [reviewed in (47)], the ability to generate distinct fibril polymorphs with unique biological properties has only recently been shown (48). We were, therefore, surprised to find that A53T PFFs generated with identical protocols using α-synuclein monomer sourced from two different companies—rPeptide and Sigma—resulted in fibrils with distinct replication capacities. While both types of PFFs induced terminal disease in TgM83^+/−^ mice inoculated via the i.c. route, only the rPeptide PFFs induced neurological signs following sc.n. injection (Figure 2). Hypothesizing that differences in titer, or specific activity, between the two fibril types could manifest as sub-clinical replication in mice inoculated with the Sigma PFFs, we used immunohistochemistry to evaluate the presence of α-synuclein neuropathology in the spinal cords of mice euthanized 475 dpi. Unlike the terminal mice inoculated with rPeptide PFFs, which exhibited robust immunostaining throughout the length of the spinal cord, mice injected with the Sigma PFFs lacked α-synuclein inclusions (Figure 4). Moreover, when we used our bioreporter cell lines to analyzed frozen brain tissue, which we previously showed are capable of detected preclinical misfolded α-synuclein (33), samples from mice injected into the sc.n. with Sigma PFFs were unable to infect cells expressing either the A53T or V55Y cell lines (Figure 5). These data suggest that while the Sigma PFFs successfully induce disease following i.c. inoculation, they are unable to efficiently replicate in the periphery.

It is not surprising that strain-specific differences in exposure route contribute to varied disease phenotypes; this is consistent with research on PrP^Sc^ prion strain biology [reviewed in (49, 50)]. For example, the hamster-adapted transmissible mink encephalopathy strains Hyper (HY) and Drowsy (DY) (6) exhibit critical differences in disease pathogenesis following inoculation via extraneural routes. Notably, while HY can neuroinvade following several routes of peripheral exposure, DY is only able to do so via a subset of these injection sites (e.g., intralingual injection is successful for transmission but intraperitoneal is not) (18, 51). Here, the unexpected finding is that the use of two different monomer sources to generate PFFs using the exact same protocol in the same laboratory resulted in two vastly different α-synuclein strains. These results, which are in alignment with similar data from So *et al*. (48), have major implications for studies that rely on PFFs to understand disease pathogenesis in synucleinopathies. The ability to generate multiple strains, even when a consistent fibrillization protocol is used, by simply altering the monomer source, means that standardization protocols, such as the protocol established via the Michael J. Fox Foundation (52), are unable to achieve aligned research tools across scientific laboratories. This finding also impacts the use of α-synuclein RT-QuIC assays as diagnostic tests (53), which are increasingly prevalent in both clinical and research use. Substantial variability and experimental outcomes between labs have been shown when using the same protocol, which may be explained by differences in monomer source (54–57). These concerns call into question the rapid proliferation of the α-synuclein RT-QuIC assay as a rigorous tool in the absence of a standardized source of α-synuclein monomer, as well as a standardized source of PFFs with a known titer to use as a positive control across laboratories.

Given our findings that the Sigma PFFs were unable to neuroinvade following sc.n. injection, we moved forward using the rPeptide A53T PFFs as the faster replicating strain in our competition studies. After first injecting TgM83^+/−^ mice with mouse-passaged MSA, a second injection with the rPeptide PFFs was performed at 30%, 45%, and 60% of the incubation period. Unexpectedly, we found that all three superinfection cohorts had a shorter incubation period than the cohort inoculated with mouse-passaged MSA only (Figure 6a), suggesting that the MSA strain was unable to interfere with PFF neuroinvasion. We, therefore, suspected that only the rPeptide PFF strain was present in the brain. Consistent with this hypothesis, brains from the terminal mice were able to infect our bioreporter cells expressing the A53T mutation, however, we then found that none of the superinfection samples were able to replicate in the V55Y cell line (Figures 6c & d). In light of our findings that the V55Y mutation exerts an inhibitory effect on this particular mouse-passage MSA patient sample (23), we interpret these results as indication that, along with the PFF strain, an inhibitory concentration of the MSA strain is present in the brains of the superinfection mice, which prevented replication in the V55Y cell line. As a result, rather than the strain competition we set out to investigate, we instead are likely seeing evidence of *strain collusion*, as the two strains appear to conspire to shorten the incubation period of disease. Excitingly, this is the first study to demonstrate the phenomenon of strain collusion, as no other co-infection study to date has described similar findings. In expanding the value of these results to all proteins that use the prion mechanism of disease, our study has identified a novel concept in prion strain biology, which warrants continued investigation.

We recognize that there are several potential limitations that may contribute to our findings. First, we previously found that during neuroinvasion of the mouse-passaged MSA strain, spinal cord pathology is not detectable via immunohistochemistry before 60% of the incubation period (23). As a result, the superinfection time points studied here could instead be forcing competition between the MSA and PFF strains to occur in the periphery rather than the spinal cord. Future studies using superinfection challenge between 60% and 100% of the incubation period would more likely discern the effect of competition in the spinal cord on disease pathogenesis. Second, differences in the underlying protein biology between PrP and α-synuclein may necessitate different approaches to study strain interference. While PrP is found on the extracellular membrane, α-synuclein is an intracellular protein found in the synapse and nucleus. As a result, differences between where the templating process occurs during neuroinvasion may impact disease pathogenesis and strain interactions, impeding our ability to use this elegant PrP^Sc^ approach to study α-synuclein strain competition. Third, simultaneous replication of both strains may lead to a collapse of the proteostasis machinery, allowing for faster propagation of both the PFF and the mouse-passaged MSA strains (58, 59).

While these are all plausible factors impacting our experimental outcomes, we hypothesize that the limitation with the greatest effect on these data is the robust overexpression of human α-synuclein in the TgM83^+/−^ mouse model. While cortical expression of the transgene is only ∼3-fold greater than endogenous α-synuclein, transgene expression in the spinal cord is almost 20-fold greater (27). In the PrP^Sc^ strain competition model, the use of non-transgenic animals limits the amount of monomeric protein available for templating (14–20). As a result, the replication efficiency, or fitness, of each strain directly impacts its ability to compete with a second strain during the self-templating process. By comparison, the remarkable overexpression of monomeric protein substrate in the TgM83^+/−^ mouse model removes the rate-limiting effect of substrate availability from impacting the interaction between the MSA and PFF strains, eliminating the need for competition. As a result, the two strains are more likely to collude with one another, rapidly spreading from the periphery into the brain. Moreover, in this scenario, a combinatorial effect of strain collusion with proteostasis collapse could further contribute to accelerated neuroinvasion. As a result, future studies that aim to understanding the potential for α-synuclein strain competition would benefit from the use of a knock-in model, removing the consequential effect of protein overexpression on experimental design and data interpretation.

In summary, the studies reported here yielded two central discoveries about α-synuclein strain biology. First, we unexpectedly found that monomer source is a critical determinant of PFF pathogenicity and replication potential. These results raise important considerations for the use of PFFs not only as a model of synucleinopathy, but also in explaining the inherent variability across α-synuclein RT-QuIC assays. This is particularly concerning as RT-QuIC-based methods become increasingly prevalent clinically as diagnostic tools. Second, we describe the first demonstration of strain collusion, where an experimental design developed to investigate strain interference instead resulted in a shortened incubation period. While a similar approach using a knock-in model, rather than overexpressing mice, may alternatively result in competition, the findings reported here raise the question of whether strain collusion could occur in individuals with duplication or triplication of the α-synuclein gene (60, 61). In these individuals, the increase in α-synuclein expression could similarly remove competition pressure, potentially allowing an α-synuclein co-pathology to replicate. It is, therefore, important to understand how distinct α-synuclein strains interact with one another, as these interactions are likely to alter the clinical course of disease.

## ACKNOWLEDGMENTS

This work was supported by grants from the NIH (R01NS121294 and accompanying supplement; R21NS127002) to A.L.W. C.R.K. is supported by a fellowship from the NIH (F31NS139652). These studies were also funded by the University of Massachusetts Amherst and Colorado State University. We thank the University of Massachusetts Amherst Animal Care Services and Colorado State University Lab Animal Resources teams for their support caring for the mice used in these studies. This work was made possible by donated human patient tissue, and we would like to recognize and thank the patients and their families for this generous gift.

## AUTHOR CONTRIBUTIONS

S.A.M.H., C.R.K., and A.L.W. designed the research; S.A.M.H., C.R.K., S.J.S., and A.L.W. performed the research; S.A.M.H., C.R.K., S.J.S., J.C.B., and A.L.W. analyzed the data; S.A.M.H., C.R.K., and A.L.W. wrote the paper; S.A.M.H., C.R.K., S.J.S., J.C.B., and A.L.W. revised the manuscript.

